# Species-wide transposable element repertoires retrace the evolutionary history of the *Saccharomyces cerevisiae* host

**DOI:** 10.1101/2021.03.01.433327

**Authors:** Claudine Bleykasten-Grosshans, Romeo Fabrizio, Anne Friedrich, Joseph Schacherer

**Affiliations:** Université de Strasbourg, CNRS, GMGM UMR 7156, Strasbourg, France; Institut Universitaire de France (IUF)

**Keywords:** Ty elements, *Ty1*, transposon activity, intraspecific variation, introgression, yeast

## Abstract

Transposable elements (TE) are an important source of genetic variation with a dynamic and content that greatly differ in a wide range of species. The origin of the intraspecific content variation is not always clear and little is known about the precise nature of it. Here, we surveyed the species-wide content of the Ty LTR-retrotransposons in a broad collection of 1,011 *Saccharomyces cerevisiae* natural isolates to understand what can stand behind the variation of the repertoire, *i.e.* the type and number of Ty elements. We have compiled an exhaustive catalog of all TE variants present in the *S. cerevisiae* species by identifying a large set of new variants. The characterization of the TE content in each isolate clearly highlighted that each subpopulation exhibits a unique and specific repertoire, retracing the evolutionary history of the species. Most interestingly, we have shown that ancient interspecific hybridization events had a major impact in the birth of new variants and therefore in the shaping of the TE repertoires. We also investigated the transpositional activity of these elements in a large set of natural isolates, and we found a broad variability related to the level of ploidy as well as the genetic background. Overall, our results pointed out that the evolution of the Ty content is deeply impacted by clade-specific events such as introgressions and therefore follows the population structure. In addition, our study lays the foundation for future investigations to better understand the transpositional regulation and more broadly the TE-host interactions.

**Authors summary:** Mobile DNA elements are widely distributed in the genomes of many eukaryotes, but their contents greatly vary between species, populations and even individuals. In fact, little is known about the origin of this variation of transposable element (TE) content across individuals of the same species. Here, we surveyed the Ty LTR-retrotransposon content in a broad collection of 1,011 *Saccharomyces cerevisiae* yeast natural isolates. We have defined an exhaustive and precise catalog of the TE variants present in the *S. cerevisiae* species. We found that the TE content follows the evolutionary history of the species because each subpopulation has a unique and specific content. Interestingly, our results highlighted that ancient interspecific hybridization events led to the appearance of new TE variants and therefore had a strong impact on the variation of the TE repertoires in this species. We also investigated the transpositional activity of these elements and found a wide variability related to the genetic background diversity. Altogether, our results have led to a better understanding of the variability of TE content at a species level.

## Introduction

Transposable elements (TEs) are interspersed repetitive DNA sequences that are able to move from one genomic location to another via a process of intra-genomic propagation, called transposition. Several decades of studies, facilitated by whole genome sequence analyzes, have shown that the genome of almost all species is populated by TEs. However, TEs are incredibly diverse, both in terms of structure and mode of transposition, as illustrated by the TE classification [1–4]. Transposable elements can be classified either into class I retrotranposons or class II DNA transposons, depending on its mechanism of transposition. In each class, TEs are grouped into orders, depending on the molecular mechanisms involved in transposition, and then divided into super-families, according to their structural characteristics. Finally, DNA sequence conservation and phylogenic data defines myriads of TE families, subfamilies and variants.

Even if TE elements are ubiquitous components, the content in terms of type, number and prevalence is very variable across the genomes. This variation was observed at different time scales, *i.e.* between species from the same phylum as well as between individuals from the same species [5–9]. In the most recent studies, the comparison of TE contents has allowed a better understanding of the events that shaped populations, such as the domestication of rice [10], silk worm [11], and sunflower [12]. Biotic or abiotic stresses have also been shown to impact the TE content of plant and insect species [13–17].

As the *Ty1* and *Ty3* elements were among the first LTR retrotransposons to be discovered, *Saccharomyces cerevisiae* emerged as a good model for TE biology [18–20]. The *Ty1* element is one of the most characterized transposons, and *Ty1* and *Ty3* have since been benchmarks for retroelement studies [21]. The in-depth knowledge of the Ty biology stands in contrast with the exploration of its species-wide diversity. This aspect remains overlooked although a large number of *S. cerevisiae* isolates have been sequenced in recent years [22–25]. In fact, *S. cerevisiae* has a seemingly simple variety of TEs with a handful of families called *Ty1* to *Ty5* [26]. All Ty elements belong to the class I and more precisely to order of LTR retrotransposons, divided into two super-families, namely copia (for the *Ty1* and *Ty2-Ty5* elements) and gypsy (for the *Ty3* elements). The common feature of their coding region is the presence of two genes similar to the *gag* and *pol* genes of retroviruses.

The Ty fraction of the *S. cerevisiae* genome is modest, around 3%, and in addition to the full-length elements, it includes a large fraction of solo-LTR resulting from the loss of the internal coding region by inter-LTR recombination. Ty elements have developed strong insertion preferences believed to generate neutral alleles [26–28]. Previous studies have already shown the variability of the Ty contents, both in terms of types and copy number [29–31]. However, these studies focused on very limited sets of isolates, preventing a global view of the evolution of the TE repertoire.

The collection of 1,011 *S. cerevisiae* natural isolates that were recently sequenced is a valuable sample to study the TE content repertoires within the species [25]. This population has a wide geographical distribution and their ecological origins are highly diverse. Its evolutionary history has delimitated 26 different subpopulations as well as three groups of mosaic isolates characterized by admixture from different lineages. In addition, a few subpopulations are characterized by introgressed regions, signatures of ancient hybridization events with the closely related species of *S. cerevisiae,* namely *Saccharomyces paradoxus* [25,32].

Here, we sought to survey the Ty content variation of *S. cerevisiae* at a species level. We first characterized all the Ty variants present in the 1,011 *S. cerevisiae* genomes by identifying new and undescribed Ty variants. We then explored the repertoire (*i.e.* the type and number of TEs) in each isolate and subpopulation. Our results clearly showed that the population structure could be defined by the variation of the Ty repertoires. In fact, each defined *S. cerevisiae* clade is characterized by its unique and specific Ty repertoire. In addition, our results highlight that introgression events had a significant impact on the appearance of new variants and therefore on the variation of repertoires between different subpopulations. Finally, we also extended our study by a functional analysis of the permissive behavior with respect to the transpositional activity, which revealed a broad variability of it across genetic backgrounds.

## Results

### Species-wide overview of the full-length Ty variants in the *S. cerevisiae* species

To explore the Ty diversity within *S. cerevisiae*, we have established an exhaustive catalog of the variants of each Ty family *(Ty1* to *Ty5)* present in the 1,011 genomes with unprecedented high resolution. To this end, the first step was to determine a set of query sequences containing the *gag-pol* coding sequences to accurately capture full-length Ty diversity. We used a set of one query for each of the five Ty families of the S288C reference genome (File S1) to perform a first BLASTn search with moderate stringency on the NCBI non-redundant nucleotide databases (Fig. S1 and see Methods). The search was conducted on both *S. cerevisiae* and *S. paradoxus* genome assemblies to obtain the most representative set of query sequences and to further resolve potential inter-species Ty flows. We have sorted hundreds of sequences based on their similarity patterns (see Methods) and obtained a final set of twelve representative queries of Ty variants distributed across the five families, adapted to cover *gag-pol* diversity across the whole *S. cerevisiae* species (File S2).

To detect already identified and even new chimeric elements in the 1,011 genomes, we performed a competitive mapping of the genomic reads on the defined set of twelve *gag-pol* sequences. We examined the coverage profiles along the queries (Fig. S1 and see Methods) and we defined a precise catalog of Ty variants found in the *S. cerevisiae* species. Out of the twelve queries, two of the *S. paradoxus* queries did not reveal significant coverage *(Ty3p* and *Ty56p).* Partial or complete coverage of the ten other queries resulted in the definition of a total of thirteen variants (Fig. 1A). Our results clearly illustrate a varying degree of diversification among the five Ty families of *S. cerevisiae* and most families do not have a large set of variants. The *Ty2* and *Ty3* families are only represented by a single variant (Fig. 1A). The *Ty4* family consists of two variants: one specific to *S. cerevisiae (Ty4c)* and another one coming from its sister species *S. paradoxus* (*Ty4p*). The *Ty5* family is also composed of two variants: a fulllength element *(Ty5f* present in a limited number of 78 isolates and a truncated version *(Ty5Δ)* present in a majority of isolates.

**Figure 1.**
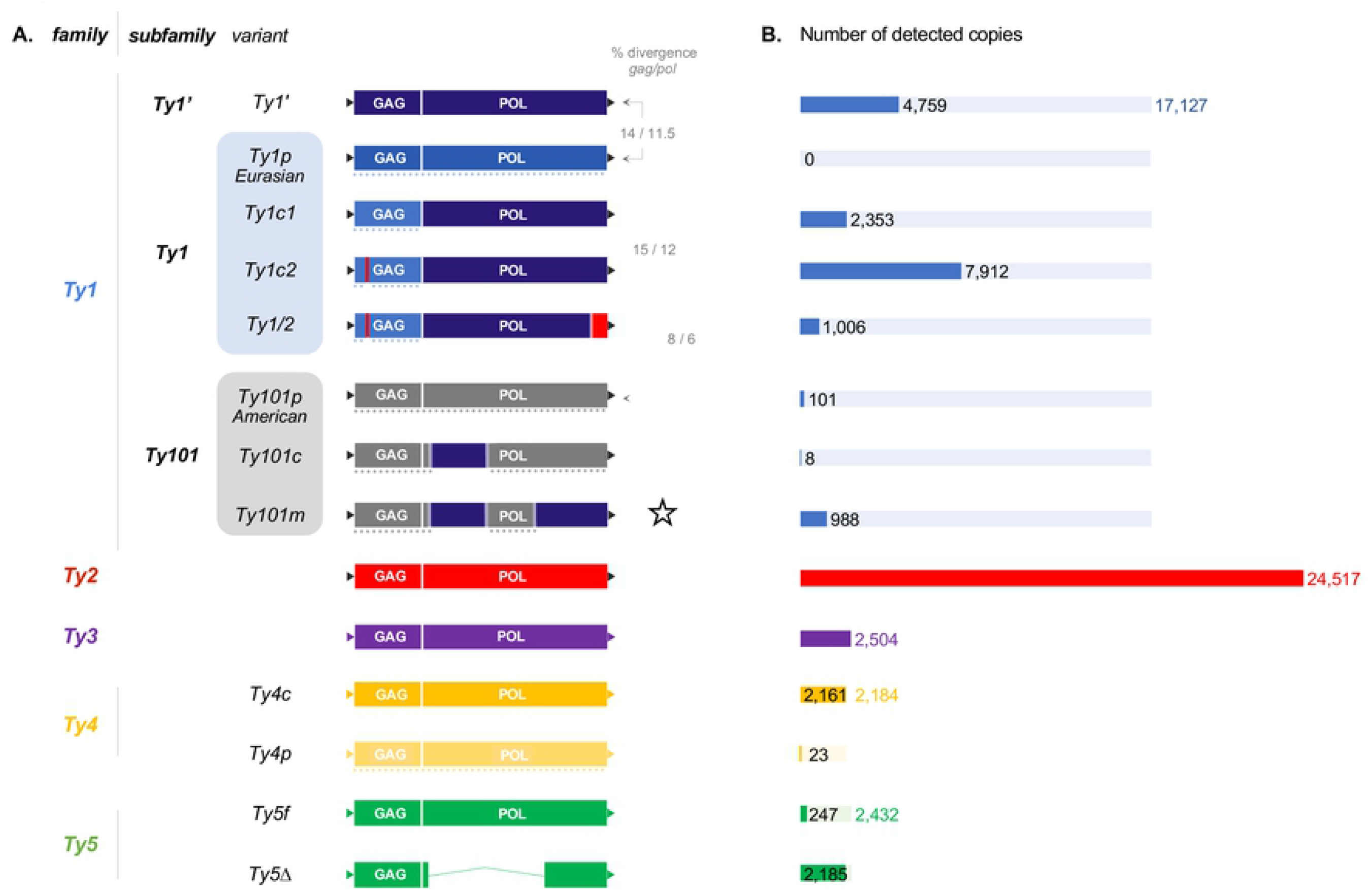
Mosaic structure and species wide prevalence of the Ty variants in *S. cerevisiae*. (**A**) The Ty variants are listed according to their classification in families and subfamilies. The color or shaded color of the Gag and Pol coding regions (boxes) and LTRs (triangles) reflect segment identity or high similarity. Dotted lines underline the Ty segments highly similar to *S. paradoxus* Ty elements. The star highlights the newly described *Ty101m* variant. The Ty elements are not represented to scale. (**B)** The size of the bars represents the total number of elements per Ty family (transparent color) and per Ty variant (solid color) detected among the 1, 011 genomes. The exact number of copies is indicated for each Ty family (colored numbers) and for each variant, if different from the family CN (black numbers).

By contrast, the *Ty1* family has a complex composition due to a high number of variants (Fig. 1A). Few of these variants have already been described [30,31,33,34] and our results provide an exhaustive catalog as well as a detailed view of their mosaic structure. Three different subfamilies can be defined based on the variability of the *gag* coding sequence. The divergence of the *gag* gene reaches 8 to 15% between the subfamilies, with the first subfamily (*Ty1*’) having a specific *gag* sequence of *S. cerevisiae* whereas the two other subfamilies *(Ty1* and *Ty101)* exhibit distinct *gag* sequences that are similar to Ty elements of *S. paradoxus.* The *gag* coding segment of the second subfamily is similar to the *Ty1p* variant and was already identified and attributed to a Eurasian *S. paradoxus* origin [31]. Here, we identified an additional interspecies transfer that involved an American lineage of *S. paradoxus*, and shaped the *Ty101* subfamily. The three subfamilies show a variable content of Ty variants. The *Ty1*’ element is the unique variant of the first *Ty1* subfamily and is composed of both *gag* and *pol* sequences specific to the *S. cerevisiae* species. The variants of the *Ty1* subfamily retain the *Ty1p gag* sequence, and they carry *pol* segments, which are overall similar to *S. cerevisiae Ty1*’. They differ only by the presence *(Ty1/2* and *Ty1c2)* or not *(Ty1c1)* of a small *Ty2* segment (Fig. 1A). The *Ty2 pol* segment in the *Ty1/2* element was already described [34] and here we have identified the *Ty1c2* element, characterized by the presence of a *Ty2* segment of 20 bp in the *gag* coding sequence. The sequence divergence between *Ty1p gag* and *Ty2 gag* reaches 28% in this segment. In the S288C reference genome, 24 out of the annotated *Ty1* elements have the *Ty1c2 gag* version while the remaining six elements have the *Ty1p gag* version. Finally, the third subfamily consists of Ty elements sharing an American *S. paradoxus gag* sequence similar to the *Ty101p* element (Fig. 1A). The *Ty101c* and *Ty101m* variants differ from *Ty101p* in that they have a mosaic *pol* sequence involving *S. cerevisiae Ty1’ pol* segments of different lengths. The *Ty101m* variant is a newly described element. We extracted the complete sequence of the *Ty101m* elements present in three isolates for which *de novo* assemblies were generated using a long-read sequencing strategy (HE015, CBS7962 and RP11.4.11) [35] and we were therefore able to confirm its mosaic structure.

### Population-scale Ty content highlights prevalence of the *Ty2* family

A total of 48,765 full-length Ty elements were detected across the 1,011 genomes (File S3). The number of Ty elements per genome is highly variable with an average of about 24 Ty elements per isolate and ranging from none to 100 elements in the PW5 and SJ5L12 isolate, respectively. The Ty families were found to be very unequal in terms of the number of elements detected (Fig. 1B). As previously shown in a small subset of strains [29,30,33,36], *Ty1* and *Ty2* are the most represented families with a total of 17,127 *(i.e.* 35.1 %) and 24,517 elements *(i.e.* 50.3 %), respectively. By contrast, the *Ty3, Ty4* and *Ty5* families are in considerably lower abundance with approximately 2,000 to 2,500 elements in each family. The distribution among isolates indicates that *Ty3* and *Ty4* are absent in a large proportion of them (absent in 511 and 593 isolates, respectively) while *Ty5* is more widely distributed (absent in 209 isolates) (File S4). *Ty3, Ty4* and *Ty5* solo-LTR were detected in all these strains (not shown), suggesting that their transpositional activity did not balance Ty loss by inter-LTR recombination.

As mentioned previously, the *Ty1* family is characterized by a large number of variants and we observed a high variability regarding their distribution. Two variants from distinct subfamilies are prevalent: the *Ty1’* and *Ty1c2* elements. While the *Ty1’* variant, specific to *S. cerevisiae,* is present in a large number of isolates (n=887) with a low copy number, the *Ty1c2* variant is the most abundant element but limited to certain isolates (n=459) (File S4). The other *Ty1* variants all show a restricted distribution in a very small subset of isolates, as well as a high degree of variability in terms of copy number. As an example, the *Ty101c, Ty101p, Ty1c1, Ty1/2* and *Ty101m* elements are present in 3 to 82 isolates with an average number of one *(Ty101c)* to 20.4 *(Ty1c1)* elements per genome.

As already mentioned, the *Ty2* element appears to be the predominant family in the *S. cerevisiae* species. Only 55 out of the 1,011 strains do not have full-length *Ty2* elements, indicating its wide distribution throughout the species. In addition, this element is also prevalent in terms of copy number with 12.7 elements per haploid genome on average (File S4). It is also present in the most divergent Asian *S. cerevisiae* isolates and, taken together, it can be considered as the ubiquitous element of the species. Regarding the origin of *Ty2*, it was reported that a very similar variant is present in some *Saccharomyces mikatae* strains but absent in the closely related *S. paradoxus* species. This observation raised the hypothesis of a horizontal transfer of *Ty2* between *S. cerevisiae* and *S. mikatae* without addressing the direction of the transfer [37]. Because *Ty2* is widespread and has a homogeneous structure, *Ty2* seems to be a recent TE element with a very active transposition in the *S. cerevisiae* species. The *Ty2* insertion polymorphism studied in isolates of various origins provides additional support for this hypothesis [30]. Nevertheless, its prevalence as well as its presence in Asian isolates strongly suggest that the *Ty2* family predates the diversification of the species [38].

### The Ty repertoires reveals the *S. cerevisiae* population structure

To have a global overview of the evolution of the Ty contents at a species-wide level, we sought to explore the conservation and variation of the Ty repertoires *(i.e.* the type and number of TEs) across the different subpopulations, which were defined based on the 1,011 sequenced genomes [25]. By clustering the strains based on the genetic diversity, we found that most of the subpopulations are defined by a unique and specific Ty repertoire (Fig 2). The generated heatmap highlights a variation of the transposon content at two levels: (i) the presence/absence of given Ty variants or elements as well as (ii) the variation in terms of copy number across the different subpopulations (Fig 2). In fact, isolates from the same subpopulation exhibit a very similar and specific repertoire, characterized by a precise type and number of TEs. These clear Ty patterns could have only been highlighted via our previous and precise characterization of the Ty elements in the large sample.

**Figure 2.**
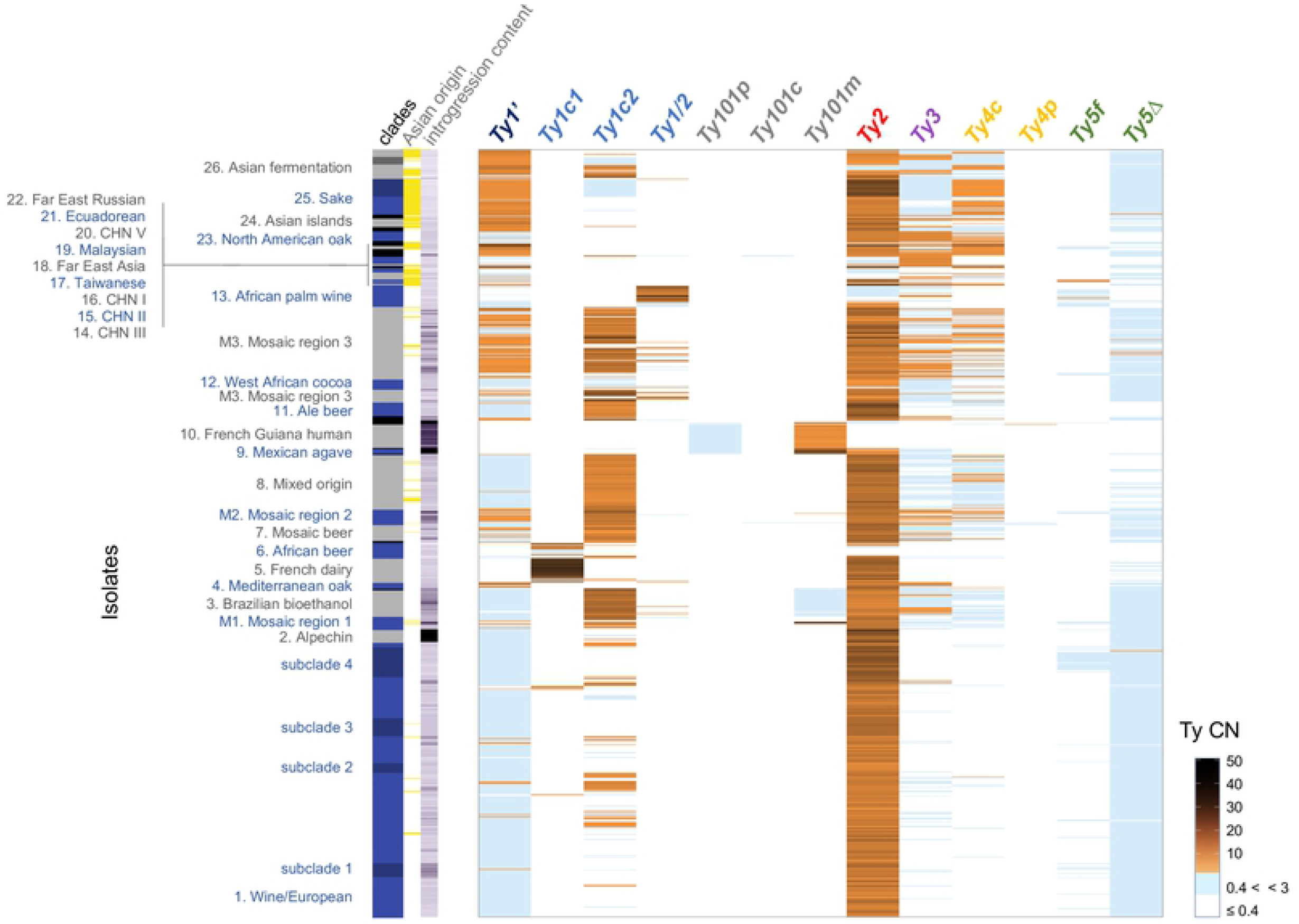
Correspondence between the Ty contents and the isolate phylogenetic relationships. The main heatmap represents the haploid-equivalent copy number of the Ty variants for each of the 1, 011 studied isolates, by using white (no Ty detected), light blue (low Ty CN < 3) and shades of orange color. The related clades are represented (left panel) by alternating of blue and gray colors. Subpopulartions are labeled darker and black is associated to strains with no clade. The yellow bars mark the isolates of Asian geographic origin. The violet shaded bars indicate the content in *S. paradoxus* ORFs ranging from 33 (light violet) to 319 (dark violet).

Our analysis shows that some Ty variants are only present in specific subpopulations, which is for example the case for the variants of the *Ty1* and *Ty101* subfamilies (Fig 2). With the exception of the *Ty1’* and *Ty1c2* variants, the other *Ty1* variants are restricted to a few subpopulations and we will detail this aspect later. Another example corresponds to the *Ty5f* variant, which is specific to the subclade 4 of the European wine as well as to the African palm wine subpopulations. Beside the clade specific TE variants, some subpopulations are also characterized by the absence of pervasive elements. As an example, the *Ty2* element is absent in the French Guiana and African beer subpopulations. By contrast, the isolates from the sake clade as well as the M1 and M2 mosaic subpopulations show a very high number of *Ty2* copies. We can also notice that the TE repertoires are characterized by specific patterns of different TE families. The most obvious example of pattern corresponds to the repertoire of the French Guiana isolates. Their Ty content is defined by the presence of *Ty101p* and *Ty101m* variants, whereas the other Ty1 variants and Ty families are completely absent. Overall, our observations emphasize the fact that the TE content is primarily shaped at the subpopulation level in *S. cerevisiae*.

Finally, the generated heatmap also highlights an interesting aspect of the evolution of the *Ty1* family (Fig 2). As mentioned previously, *Ty1’* is the *S. cerevisiae* specific and prevalent element of the *Ty1* family. *Ty1’* segments are almost ubiquitous across the 1,011 isolates as they are present in the mosaic *Ty1* variants, such as the variants from the *Ty1* and *Ty101* subfamilies. Nevertheless, we observed an enrichment of the full-length *Ty1’* variant in some specific subpopulations and more precisely in the Asian isolates (Fig S3). This is an interesting observation in respect to the single ‘out of China’ origin of the *S. cerevisiae* species [25]. The geographic distribution as well as its omnipresence provide a strong support to the fact that *Ty1’* is the ancestral representative element of the *Ty1* family, as recently hypothesized [31].

### Contrasting outcomes of independent interspecies hybridization events on the Ty contents

Our population genomic study focusing on the 1,011 *S. cerevisiae* genomes highlighted the presence of introgressed ORFs (Open Reading Frames) coming from the closely related species, *S. paradoxus* [25]. In fact, all studied isolates carry at least one introgressed ORF, with a mean of 32, indicating a ubiquitous gene flow between these yeast species. In this context, it was interesting to find a large number of TE variants of the *Ty1* family, being chimeric elements between *S. cerevisiae* and *S. paradoxus*. As previously described, the *Ty1* diversity was shaped by introgression events with Eurasian and American *S. paradoxus* lineages, leading to the *Ty1* and *Ty101* subfamilies, respectively (Fig 2). These introgression events correspond to secondary contacts with *S. paradoxus* and occurred after the out-of-China dispersal [25]. They consequently had a major impact on the genome evolution in terms of gene content (introgressed ORFs) but also clearly shaped the Ty repertoire at the subpopulation level (Fig 2). Whereas the *Ty1’* variant is probably the ancestral element of this subfamily, the *Ty1* and *Ty101* variants are the result of Ty flow between these sister species.

Regarding the introgression events involving the American *S. paradoxus* lineage, the Mexican agave and French Guiana subpopulations were found to have a large number of introgressed ORFs (n=207 and n=86 on average, respectively). These two subpopulations were impacted by this major event as they retained copies of the *S. paradoxus Ty1* element *(Ty101p)* and contain mosaic elements *(Ty101m)* (Fig 3). Whereas the two subpopulations probably lost the *Ty1’* element, only the Mexican agave subpopulation kept the *Ty2* element, leading to a slightly different repertoire. Finally, we had the opportunity to determine the genomic location of the *Ty101m* elements in three isolates (HE015, CBS7962 and RP11.4.11) for which the genomes were completely assembled using Oxford Nanopore sequencing [35]. By comparing the location of the introgressed genes and the TEs, it was interesting to observe that there is no overlap between the *Ty101m* elements and this set of genes. This observation provides an evidence of the existence of active *Ty101m* elements.

**Figure 3.**
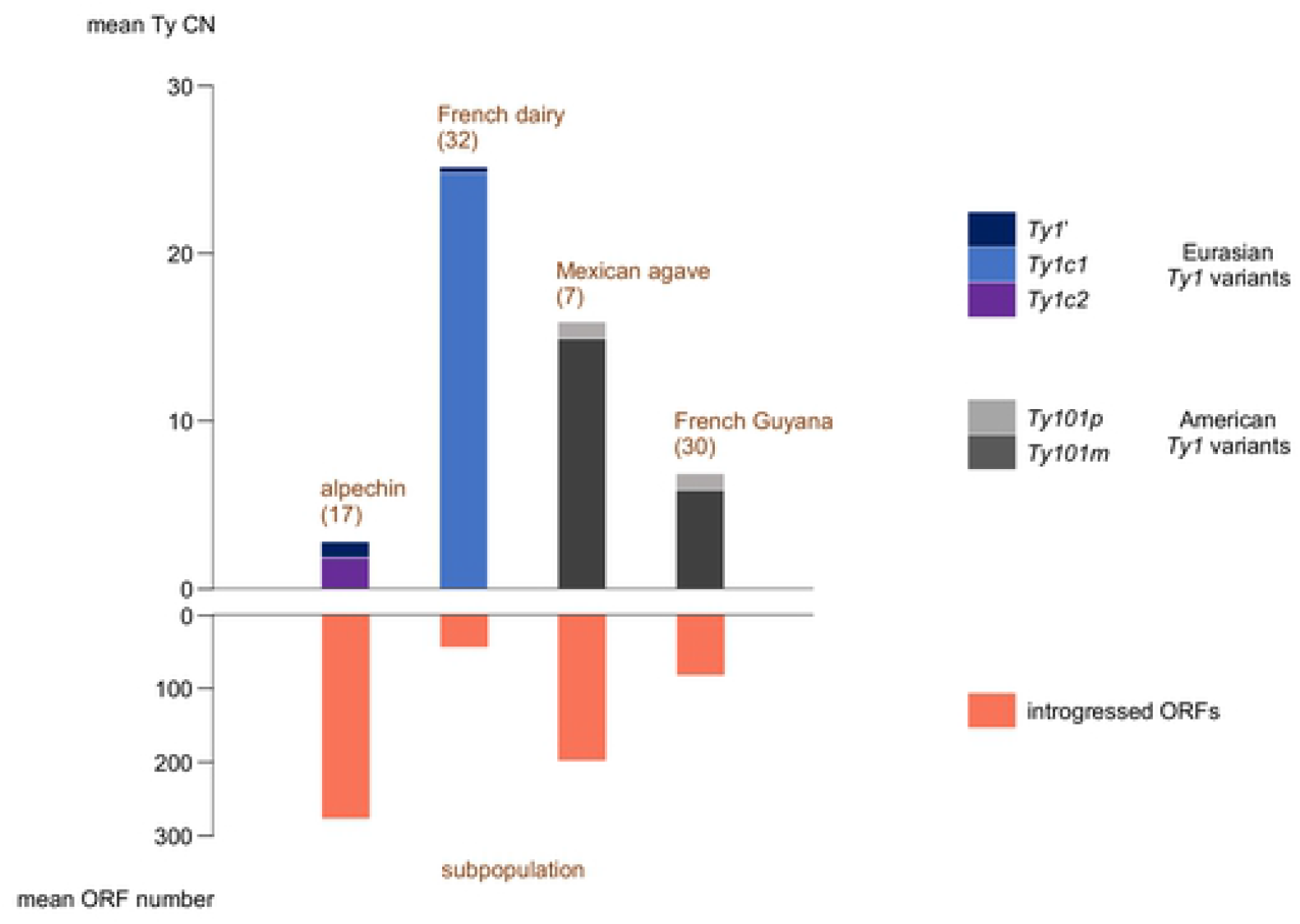
Outcomes of ancestral hybridization in terms of *Ty1* and introgressed ORF contents in four subpopulations. In the bar-plot, the upper bars represent the mean number of the *Ty1* variants. The lower bars represent the number of introgressed *S. paradoxus* ORFs. The number of isolates in the subpopulations is indicated in brackets.

The introgression events involving the Eurasian *S. paradoxus* lineage led to a contrasting evolution of the Ty repertoires. The Alpechin subpopulation, with the highest number of introgressed ORFs (n=287 on average), exhibit a Ty landscape with almost no *S. cerevisiae* / *S. paradoxus* mosaic elements (Fig 3). Only five out the 17 isolates carry some *Ty1c2* elements. We then analyzed the Ty content of the hybrid ancestor of the Alpechin subpopulation, which was recently described [32]. By using our set of Ty queries, we detected six *Ty1p* elements in this genome. This observation indicates that the *S. paradoxus Ty1p* is present in this ecological niche and was present in the founder population of the current *S. cerevisiae* Alpechin subpopulation. It is therefore clear that neither this original element nor a mosaic version of it did play a major role in shaping the Ty landscape. By contrast, the French dairy clade with a small of introgressed ORFs (n=48 on average) is characterized by the presence of a high number of *Ty1c1* elements. This variant is almost exclusive to this subpopulation and is only found in a few isolates, such as the African beer strains.

Altogether, these results highlight the clade-specific and differential impacts of the introgression events on the Ty repertoire. They also revealed the origin of the broad diversity of *Ty1* variants. With the exception of the *Ty1c2* variant, the other variants are mostly private or specific to few subpopulations. This high diversity of *Ty1* variants stands in contrast with the low number of variants present in the other subfamilies. Even if the *Ty2* is almost ubiquitous, only one variant is detected in our large population, probably as a consequence of the absence of *Ty2* in *S. paradoxus*.

### Variable transpositional activity across genetic backgrounds

Beside the Ty content diversity, we also sought to address transpositional activity variation across this natural population. In fact, the evaluation of permissive or restrictive transposition behaviors with respect to the Ty content could shed light on their mobilization. With this objective in mind, we determined the transpositional activity in a large set of 92 natural isolates as well as 79 stable haploid derivatives (File S5). To determine the transpositional activity, previous studies used a reporter toolbox based on an element, called *Ty1his3-AI,* that carries an antisense inactive *his3-AI* allele disrupted by an artificial intron [39]. The reverse-transcription of this element based on the transposition process can restore a functional *HIS3* allele after the *AI* intron splicing. This system is commonly used in haploid strains carrying a resident *his3* defective allele [21].To assess the transpositional activity of natural isolates with variable ploidy levels, we constructed and validated a new reporter that allows a direct selection (see Methods and File S6). This reporter, called *Ty1hygro-AI,* carries the same *Ty1* variant than the original construction [31,40] but the *his3-AI* cassette was replaced by a *hygro-AI* cassette, allowing a direct selection without any genetic manipulation of the natural isolates (see Material and Methods).

We then used this construction to investigate the transpositional activity of 92 natural isolates, which were chosen to cover the broad genetic diversity of the *S. cerevisiae* species. We found that the frequency of the Ty activity across the natural isolates is variable, ranging from 0.45 to 273 × 10^−6^ (File S5 and Fig 4B). In addition, we also investigated a collection of 79 generated stable haploid strains [41]. For the set of haploid strains, we found that the frequency of the Ty activity exhibits a higher variability, ranging from 0.35 to 1870 × 10^−6^ (File S5 and Fig 4A and B). The distribution of the activity is bimodal with two groups including a set of strains (n=20) that display a strong restrictive behavior toward transposition and another one (n=40) encompassing permissive isolates.

**Figure 4.**
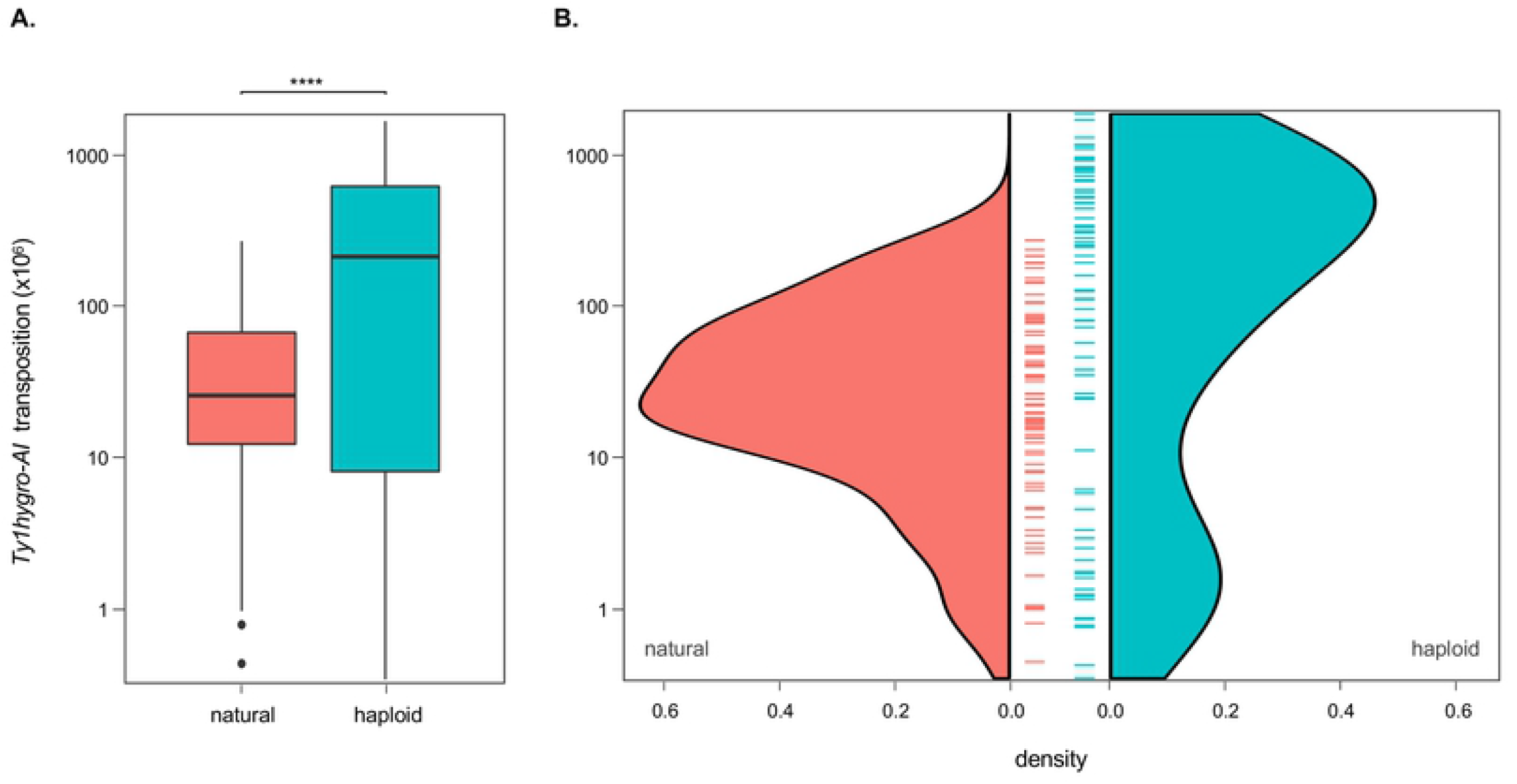
Species wide variation in the *Ty1hygro-AI* transpositional activity. The transpositional activity represented as the ratio of the hygromycine-resistant colonies to the total cells was assessed in 93 natural isolates and in 79 haploid strains. Each value is the mean of triplicate. The distribution, the median (**A**) and the probability density function (**B**) are represented.

All these strains were also selected to lead to a significant overlap between the sets of haploid and natural isolates. Consequently, we compared the transpositional activity at two ploidy levels (n and 2n) for a total of 45 genetic backgrounds (Fig S5). Our results show an overall higher transpositional activity in the haploid compared to the diploid isolates (Fig 4A, p-value = 2.1 × 10^−5^, Wilcoxon). This is in agreement with previous studies where it was found that *Ty1* transcription is repressed in a *MATa/MATα* diploid context [42,44]. Even if this observation is a general trend, we still observe a variability across the different genetic backgrounds. Interestingly, most of the strongly restrictive haploid strains (n=10) display a lower transpositional activity compared to their corresponding diploid (Fig S5). This observation therefore highlights that *Ty1* transposition repression in diploids cannot be generalized and is dependent of the genetic background.

Finally, we explored the correlation between the transpositional activity and the Ty contents for both the haploid and the natural isolates. Our results show an absence of correlation between the Ty activity and the number of copies of the main families or subfamilies *(Ty1, Ty1’* and *Ty2)* in both sets of strains (Fig S6). This observation is somewhat at odds with previous studies showing that the *Ty1* activity is decreased when additional copies of the *Ty1* element are present in the genome, a mechanism termed Copy Number Control (CNC) [31,45–47]. If this mechanism would have been a predominant mechanism acting on pervasive transposition at a population-scale, we would expect to observe an anti-correlation between the Ty content and activity for such a large sample. CNC has been studied in a very limited number of isolates and this mechanism might be genetic background specific. In addition, we cannot exclude that CNC may be a preponderant response of the host during a burst of transposition but that additional regulation mechanisms might act in isolates already carrying *Ty1* elements.

## Conclusion

To investigate the composition of the repetitive genomic fraction represented by the full-length Ty retrotransposons, we developed a strategy to dissect this aspect across the whole genome sequences of 1,011 *S. cerevisiae* isolates. We first determined the catalog of *Ty* variants present in this species with an unprecedented level of precision. Our results highlighted that the Ty landscape retraces the population structure with each subpopulation having a unique and specific TE repertoire.

Among the five Ty families (*Ty1* to *Ty 5*), we have shown that the *Ty1* family is the more diversified one as three subfamilies were identified, each including one to four variants. Whereas the *Ty1’* variant is most likely the ancestral element, the *Ty1* and *Ty101* subfamilies are the results of subpopulation events, namely hybridizations with distinct lineages of the *S. paradoxus* species. Interestingly, *Ty2* which is the preponderant Ty family does not present such a diversification, probably due to the fact that there is no *Ty2* elements in *S. paradoxus.* Overall, our results revealed the impact of the interspecific hybridization events on the evolution of the Ty landscape.

The diversification of the panoply of Ty elements in *S. cerevisiae* comes from the generation of mosaic elements that combine segments originating from several species-specific TEs. The process that can lead to such mosaic elements is known as inter-element recombination and it was already proposed to explain the existence of the *Ty1/2* hybrids [34]. While ectopic homologous recombination between Ty copies can occur [48–50], these mosaic elements can also be the result of the reverse-transcriptase jumping between distinct RNA templates that are encapsidated in the same virus-like particle, like it was described for recombinant retroviruses [51–53]. The latest mechanism was already shown to be highly efficient in the generation of *Ty1* sequence diversity in the S288C reference strain [54].

Overall, it is of particular interest to see the rise of new Ty repertoires generated by the successful interspecific hybridization of distant genomes and associated Ty elements. By contrast to the case of hybrid dysgenesis syndrome in some *Drosophila* species [55], it is not possible to predict if these TEs will lead or not to a genomic shock [56] and detrimental TE proliferation that can ultimately result in reproductive isolation. Furthermore, there is growing evidence that hybridization events and their associated TE contents is not obligatory subjected to massive TE deregulation. Such outcomes were described in yeast species [57,58], as well as in other groups including plants [59,60] and insects [61,62]. Our results show that these hybridization events can lead to different and unpredictable evolutionary trajectories as well as Ty repertoires. Interestingly, hybridization events can lead to new TE variants and this reveals some possible long-term impact of natural hybridization on the TE landscape. By analogy with the transgressive phenotypic traits that arise through hybridization [63,64], these new TE repertoires can therefore being shaped during the so-called reticulated evolution [65]. Exploration of species that result from hybridization events, exhibit admixed populations or contain recursive hybrids [66] will therefore be insightful to dissect the formation, the evolution and the impact of these newly emerging TE repertoires.

## Materials and methods

### Yeast strains

The 1,011 isolates investigated for Ty content were described in [25] and are listed in File S3. The strains and isolates used for the transposition assays are listed in File S5.

### Set-up of the *gag-pol* query collection (Figure S1)

BLASTn searches were conducted on the NCBI non-redundant nucleotide database (July 6, 2018), to construct a collection of *gag-pol* sequences from the *S. cerevisiae* and *S. paradoxus* species. We have used a *gag-pol* sequence from the S288c reference genome, for each of the *Ty1* to *Ty4* family, as query sequences (File S1). As the single *Ty5* copy in S288c carries a 1500 bp internal deletion, we included the sequence of the full-length *Ty5* from YJM178 strain. Target sequences with complete coverage along each query were sampled and aligned (Clustal Omega, https://www.ebi.ac.uk/), exact duplicates were excluded. Among sequences sharing more than 95 % identity, a single representative was retained. Additional manual inspections of the alignments were performed, leaving a set of twelve query sequences (File S2) that we considered representative of the Ty diversity in both *S. cerevisiae* and *S. paradoxus* species. This set of *gag-pol* query sequences was used to detect Ty elements among the 1,011 genomes.

### Detection of sequence reads by mapping on Ty segments

Reads mapping was performed with bwa [67] using the previously described set of *gag-pol* sequences as reference. Samtools mpileup [68] was used to generate pileup files that were processed to estimate the coverage along the reference sequences (File S2) using 10 bp non-overlapping windows.

### Assignation to Ty families or variants and estimation of the Ty copy number

Coverage profiles were obtained by plotting the reads depth along the corresponding *gag-pol* Ty reference (examples in Fig S2). To address the mosaic organization of Ty variants and detect new mosaic variants, the coverage plots were manually inspected. This allowed to define (i) the appropriate segments for accurately assigning the detected elements to a Ty category (family, subfamily or variant), and (ii) the most appropriate region to assess CN from the reference coverage. A summary of the considered regions is provided hereafter:

- *Ty1:* region from 100 to 1,000 of the *Ty1’* query, that corresponds to the *gag* segment and differentiates the *Ty1* subfamilies;
- *Ty1c2:* region from 100 to 1,000 of the *Ty1c2* query;
- *Ty2*: region from 100 to 4,500 of the *Ty2* query;
- *Ty3c/p* and *Ty4c/p:* the whole respective query;
- *Ty5f:* region from 1,300-3,400 of the *Ty5* query. *Ty5Δ* was obtained by the difference of coverage between the 3,450-4,800 and the 1,300-3,400 segments on the *Ty5* query.

The *Ty1/2* elements deserved specific attention. Indeed, they were characterized by an excess of coverage at the 4,900-5,100 region of the *Ty2* query, balanced by a drop in the coverage of the 4,900-5,300 region of the *Ty1c2* query (Fig S2 A, strain S288C). If only *Ty1/2* are present *(i.e.* the coverage on the *Ty1c2* 4,900-5,300 segment resulted in a value less than 0.5) we inferred the CN computed on the 100-1,100 segment of the *Ty1c2* query as the actual *Ty1/2* CN. When both *Ty1* and *Ty1/2* are present, the *Ty1/2* CN was assessed by the difference of coverage between the *Ty2* 4,900-5,100 and 100-4,500 segments. The corresponding value was then subtracted to the value computed on the 100-1,100 segment of the *Ty1c2* query, in order to adjust the *Ty1c2* CN.

Among the *Ty1* family, the following variants were carefully considered:

- *Ty1c1* differs from *Ty1c2* by a very short segment in the *gag* region. *Ty1c1* was revealed by an excess of coverage of this short segment on the *Ty1p* query (Fig S2 A, strain S288C). If *Ty1c1* are the only *Ty1* representatives within a strain, CN was inferred from the coverage of the 100-1,100 region of the *Ty1c2* query. Otherwise, they were recorded by default as *Ty1c2* elements;
- *Ty101m* variant was revealed by balanced coverage and absence of coverage of adjacent segments of the *Ty101c, Ty101p* and *Ty1’* reference sequences (Fig S2 A, strain HE015). CN was computed from the coverage along the *Ty101c* 500-1,000 segment;
- *Ty101c* and *Ty101p* variants were detected by manual inspection of the coverage plots and their CN computed from the coverage along the 500-1,000 segment of the corresponding query.

Finally, the Ty CN related to haploid-equivalent genome was estimated as the ratio between the average of the computed coverage and the whole nuclear genome coverage [25]. Assessments for haploid-equivalent genomes enable further comparisons between isolates of variable ploidy levels (File S3).

### LTR detection

We aimed to detect the presence/absence of the Ty LTR, in order to identify isolates where solo-LTRs are the sole representatives of a given a Ty family. We used a set of queries including three LTR sequences that are representative of the very similar *Ty1* and *Ty2* LTRs and one LTR sequence for each of the *Ty3* to *Ty5* families (File S7).

### Yeast methods

Yeast cells were grown on YPD (yeast extract 1%, peptone 2%, dextrose 2 %) at 30°C. When required, the temperature was adjusted to 34°C or to 22°C which are respectively restrictive or permissive to *Ty1* transposition [69,70]. Geneticin (Euromedex) to a final concentration of 200 μg/ml was added to obtain the selective YPD-G418 media. The solid selective media YPD-hygro was supplemented with hygromycin (Euromedex) to a final concentration of 200 μg/ml and with 2% agar (Bacto agar Difco). Composition of the nitrogen-depleted media was as follows: 0.67% Yeast Nitrogen Base without amino acids and ammonium sulfate (MP Biomedicals), 2% D-glucose, 0.05 mM ammonium sulfate. Yeast strains were transformed by the *Ty1hygro-AI* plasmid with the EZ-Yeast Transformation Kit (MP Biomedicals).

### *Ty1hygro-AI* plasmid construction and validation

The pCeTyX *Ty1hygro-AI* carrying plasmid was constructed by assembling four overlapping DNA fragments thanks to the Gibson Assembly Cloning Kit (NEB). The 5,472 bp plasmid backbone was obtained after restriction of p41Neo 1-F GW (https://www.addgene.org/58545/) by *XhoI* and *XbaI.* We designed and purchased the *hygro-AI* cassette at Genescript and further amplified it by PCR using Iproof high fidelity DNA polymerase (BioRad). The 5’-LTR-*gagpol* and the 3’-LTR segments were obtained by PCR amplification from the pOY1 template [40]. The sequence of the final construction was verified by Sanger sequencing (Eurofins). Details on pCeTyX cloning and primer sequences are available in File S6.

We first validated the new *Ty1hygro-AI* reporter by testing its activity in different conditions in the S288C genetic background. The results obtained with the *Ty1hygro-AI* reporter recapitulates the available results of *Ty1his3-AI* reporters, as illustrated in (Fig S4 A). Transposition was observed to be higher in the haploid state (FY5) compared to the diploid state (FY3) [43], optimally at 22°C. Further the transposition activity was undetectable at 34°C [70]. We also have recorded the transpositional activity of the *Ty1hygro-AI* reporter in the CLQCA_20-259 background (File S5), where no Copia elements were detected. Importantly, this shows that the *Ty1hygro-AI* reporter is autonomous.

### *Tylhygro-AI* mobility assay

To reliably assess the *Ty1hygro-AI* transpositional activity in a large number of strains, we designed and performed the following protocol in triplicate. Strains were transformed with pCeTyX, and bulk G418 resistant transformants were selected in 1.8 ml YPD-G418, at 34°C (restrictive temperature for *Ty1* transposition) during 48 h. Subsequently, 10 μl of each culture were transferred to 150 μl fresh YPD-G418 and grown to saturation for 36 h at 34°C. To allow for *Ty1* transposition, 100 μl of the saturated cultures were inoculated in 1.8 ml YPD and incubated at the permissive temperature of 22°C for 24h. Variable fractions ranging from 0.1 to 95 % of the cells were harvested, plated on solid YPD-hygro, and incubated for 48 to 72 h at 34°C. Hygromycine-resistant colonies were counted. For each assay, a 50 μl aliquot of the induced cultures was saved for further cell counting. Cell counting was performed by counting of the events with FCS ≥ 1.10^4^ and SSC ≥ 5.10^5^ by flow cytometry (BD-Accuri-C6). The transposition activity was represented as the ratio of the hygromycine-resistant colonies to the total cells. Depending on the efficiency of the transformation step, this protocol ensures that at least three independent transformants were assessed for each of the strains. In parallel, to control for the absence of preexisting transposition events that might have occurred prior to the induction step at 22°C, 20 μl of the aforementioned saturated cultures at 34°C were deposited on solid YPD-hygro and incubated during 48 h at 34°C; if hygromycine-resistant colonies grew on this control, the corresponding assay was discarded.

Some genetic backgrounds display very low ratio of hygromycine-resistant colonies in the standard assay conditions. In order to rule out issues with the assay, two additional controls were performed. In the first control, we subjected the corresponding strains to conditions of nitrogen starvation during the step of *Ty1* induction. Nitrogen starvation is known to stimulate the *Ty1* transposition (Fig S4 A and [43]) and all the tested strains show substantial increases of the ratio in hygromycine-resistant colonies in this condition (examples in Fig S4 B). This indicates that the *Ty1hygro-AI* reporter is actually functional even for the strains that display low transpositional activity very close to the detection threshold in the usual assay conditions. In the second control, we checked if pCeTyX may be lost during the step of *Ty1* induction. This control was performed in several strains with variable transpositional activity. The results indicate that at least 40 % of the cells retained pCeTyX after 24 h of growth at 22°C in non-selective media. The tested strains display variable rates of pCeTyX plasmid loss, however the strains with the lowest activity have the highest rates of plasmid retention and *vice-versa* (Fig S4 C).

## Acknowledgments

We thank Joan Curcio for the pOY1 plasmid. We also thank members of the Schacherer lab at the University of Strasbourg for helpful suggestions throughout the project.

## Supplemental Figures

**Supplemental Figure 1. Main steps of the strategy used to characterize the 1,011 genome Ty contents.**

**Supplemental Figure 2. Coverage based Ty CN computation.** The correspondence between the coverage profiles along nine Ty queries (**A**) and the Ty contents (**B**) is illustrated for the S288C reference strain and the HE015 isolate. The query coverages are normalized to the genome coverage of each strain, the two horizontal dashed lines indicate 1x and 10x levels. The profiles along *Ty3p*, *Ty4p* and *Ty56p* queries are not represented due to the fact that they are empty for the two strains. The Ty CN par haploid genome computed according to representative query segments specific for each Ty variant (this study) are compared to alternative CN determinations.

**Supplemental Figure 3. Enrichment of the *Ty1’* subfamily in isolates of Asian geographical origin.** Pairwise comparisons were performed using Wilcoxon rank sum test with Benjamini-Hochberg adjustment for multiple comparisons.

**Supplemental Figure 4. Validation of the *Ty1hygro-AI* reporter.** (**A**) Squared areas of the same YPD-hygro plate were spread with approx. 5.10^8^ cells of FY5 or FY3 strains carrying pCeTyX and subjected to 24 hours of growth in different conditions. Cell numbers were adjusted after OD600 measurements. The plate was incubated at 34°C during 72 hours. (**B**) YPD-hygro plate areas were spread with approx. 1.10^8^ cells of different genetic backgrounds carrying pCeTyX and subjected to 24 hours of growth at 22°C in YPD or in nitrogen-depleted media, and incubated at 34°C during 72 hours. Cell numbers were adjusted after OD600 measurements. (**C**) Cells of different genetic backgrounds carrying pCeTyX were subjected to growth on non-selective YPD during 24 hours and serial dilutions were spread on solid YPD. After 48 hours at 30°C, colonies were replicated on solid YPD-G418. pCeTyX retention was estimated as the ratio of G418-resistant to total colonies. In parallel, the transpositional activity of the samples was assessed according to the standard assay protocol.

**Supplemental Figure 5. Comparison of *Ty1hygro-AI* activity between 45 natural isolates and the corresponding haploid strain.** The transpositional activity in the diploid state is plotted as a function of the transpositional activity in the haploid state. The lighter blue points correspond to the values of the isolates that display very low transpositional activity in the haploid corresponding strains. The dotted line indicates the position of the linear function f(x)=x.

**Supplemental Figure 6. Absence of correlation between Ty CN and *Ty1hygro-AI* transpositional activity.** The transpositional activity is plotted as a function of the main Ty variant CN. Blue and red colors are used for the haploid and diploid values, respectively. The lighter blue points correspond to the values of twelve isolates that display very low transpositional activity in the haploid corresponding strains. Kendall’s rank correlation coefficients are indicated.

## Supplemental Files

**Supplemental File 1. Sequence and reference of the five basic Ty queries.**

**Supplemental File 2. Sequence and reference of the twelve representative Ty queries.**

**Supplemental File 3. Copy number of the Ty variants in the 1, 011 *S. cerevisiae* isolates.** The copy number are computed for haploid equivalent genomes. The isolates are listed according to their position in the neighbor-joining tree of the species established in [25].

**Supplemental File 4. General statistics of Ty families and variants.** Number and fraction (%) of detected Ty elements; mean and median Ty number per haploid equivalent genome; number of carrying isolates, mean Ty number in carrying isolates (haploid equivalent).

**Supplemental File 5. *Ty1hygro-AI* transpositional activity for 92 natural isolates and 79 haploid strains.** The activity is represented by the ratio between hygromycin resistant and total cells resulting from independent triplicates. The stimulation of the transpositional activity in nitrogen-depleted media is indicated for the strains and isolates for which it was tested. The compared activities in the overlapping set of haploid strains and natural isolates are summarized in a separate sheet.

**Supplemental File 6. pCeTyX cloning: information on assembled fragments and primer sequences.**

**Supplemental File 7. Sequence and reference of the six LTR queries.**

## References

1. Wicker T, Sabot F, Hua-Van A, Bennetzen JL, Capy P, Chalhoub B, et al. A unified classification system for eukaryotic transposable elements. Nature Reviews Genetics. 2007;8: 973–982. doi:10.1038/nrg2165

2. Arkhipova IR. Using bioinformatic and phylogenetic approaches to classify transposable elements and understand their complex evolutionary histories. Mobile DNA. 2017;8: 19. doi:10.1186/s13100-017-0103-2

3. Bourque G, Burns KH, Gehring M, Gorbunova V, Seluanov A, Hammell M, et al. Ten things you should know about transposable elements. Genome Biology. 2018;19: 199. doi:10.1186/s13059-018-1577-z

4. Kojima KK. Structural and sequence diversity of eukaryotic transposable elements. Genes & Genetic Systems. 2019;94: 233–252. doi:10.1266/ggs.18-00024

5. Huang CRL, Burns KH, Boeke JD. Active Transposition in Genomes. Annual Review of Genetics. 2012;46: 651–675. doi:10.1146/annurev-genet-110711-155616

6. Thomas-Bulle C, Piednoël M, Donnart T, Filée J, Jollivet D, Bonnivard É. Mollusc genomes reveal variability in patterns of LTR-retrotransposons dynamics. BMC Genomics. 2018;19: 821. doi:10.1186/s12864-018-5200-1

7. Gaiero P, Vaio M, Peters SA, Schranz ME, de Jong H, Speranza PR. Comparative analysis of repetitive sequences among species from the potato and the tomato clades. Ann Bot. 2019;123: 521–532. doi:10.1093/aob/mcy186

8. Ray DA, Grimshaw JR, Halsey MK, Korstian JM, Osmanski AB, Sullivan KAM, et al. Simultaneous TE Analysis of 19 Heliconiine Butterflies Yields Novel Insights into Rapid TE-Based Genome Diversification and Multiple SINE Births and Deaths. Genome Biol Evol. 2019;11: 2162–2177. doi:10.1093/gbe/evz125

9. Fonseca PM, Moura RD, Wallau GL, Loreto ELS. The mobilome of Drosophila incompta, a flower-breeding species: comparison of transposable element landscapes among generalist and specialist flies. Chromosome Res. 2019;27: 203–219. doi:10.1007/s10577-019-09609-x

10. Carpentier M-C, Manfroi E, Wei F-J, Wu H-P, Lasserre E, Llauro C, et al. Retrotranspositional landscape of Asian rice revealed by 3000 genomes. Nature Communications. 2019;10: 24. doi:10.1038/s41467-018-07974-5

11. Han M-J, Xu H-E, Xiong X-M, Zhang H-H. Evolutionary dynamics of transposable elements during silkworm domestication. Genes Genom. 2018;40: 1041–1051. doi:10.1007/s13258-018-0713-1

12. Mascagni F, Vangelisti A, Giordani T, Cavallini A, Natali L. Specific LTR-Retrotransposons Show Copy Number Variations between Wild and Cultivated Sunflowers. Genes. 2018;9: 433. doi:10.3390/genes9090433

13. Quadrana L, Bortolini Silveira A, Mayhew GF, LeBlanc C, Martienssen RA, Jeddeloh JA, et al. The Arabidopsis thaliana mobilome and its impact at the species level. Zilberman D, editor. eLife. 2016;5: e15716. doi:10.7554/eLife.15716

14. Stritt C, Gordon SP, Wicker T, Vogel JP, Roulin AC. Recent Activity in Expanding Populations and Purifying Selection Have Shaped Transposable Element Landscapes across Natural Accessions of the Mediterranean Grass Brachypodium distachyon. Genome Biol Evol. 2018;10: 304–318. doi:10.1093/gbe/evx276

15. Lerat E, Goubert C, Guirao-Rico S, Merenciano M, Dufour A-B, Vieira C, et al. Population-specific dynamics and selection patterns of transposable element insertions in European natural populations. Molecular Ecology. 2019;28: 1506–1522. doi:10.1111/mec.14963

16. Rogivue A, Choudhury RR, Zoller S, Joost S, Felber F, Kasser M, et al. Genome-wide variation in nucleotides and retrotransposons in alpine populations of Arabis alpina (Brassicaceae). Molecular Ecology Resources. 2019;19: 773–787. doi:10.1111/1755-0998.12991

17. Baduel P, Quadrana L, Hunter B, Bomblies K, Colot V. Relaxed purifying selection in autopolyploids drives transposable element over-accumulation which provides variants for local adaptation. Nature Communications. 2019;10: 5818. doi:10.1038/s41467-019-13730-0

18. Cameron JR, Loh EY, Davis RW. Evidence for transposition of dispersed repetitive DNA families in yeast. Cell. 1979;16: 739–751. doi:10.1016/0092-8674(79)90090-4

19. Clark DJ, Bilanchone VW, Haywood LJ, Dildine SL, Sandmeyer SB. A yeast sigma composite element, TY3, has properties of a retrotransposon. J Biol Chem. 1988;263: 1413–1423.

20. Hansen LJ, Chalker DL, Sandmeyer SB. Ty3, a yeast retrotransposon associated with tRNA genes, has homology to animal retroviruses. Mol Cell Biol. 1988;8: 5245–5256. doi:10.1128/mcb.8.12.5245

21. Curcio MJ, Lutz S, Lesage P. The Ty1 LTR-retrotransposon of budding yeast, Saccharomyces cerevisiae. Microbiol Spectr. 2015;3: 1–35. doi:10.1128/microbiolspec.MDNA3-0053-2014

22. Strope PK, Skelly DA, Kozmin SG, Mahadevan G, Stone EA, Magwene PM, et al. The 100-genomes strains, an S. cerevisiae resource that illuminates its natural phenotypic and genotypic variation and emergence as an opportunistic pathogen. Genome Res. 2015;25: 762–774. doi:10.1101/gr.185538.114

23. Zhu YO, Sherlock G, Petrov DA. Whole Genome Analysis of 132 Clinical Saccharomyces cerevisiae Strains Reveals Extensive Ploidy Variation. G3 (Bethesda). 2016;6: 2421–2434. doi:10.1534/g3.116.029397

24. Gallone B, Steensels J, Prahl T, Soriaga L, Saels V, Herrera-Malaver B, et al. Domestication and Divergence of Saccharomyces cerevisiae Beer Yeasts. Cell. 2016;166: 1397–1410.e16. doi:10.1016/j.cell.2016.08.020

25. Peter J, De Chiara M, Friedrich A, Yue J-X, Pflieger D, Bergström A, et al. Genome evolution across 1,011 Saccharomyces cerevisiae isolates. Nature. 2018;556: 339–344. doi:10.1038/s41586-018-0030-5

26. Lesage P, Todeschini AL. Happy together: the life and times of Ty retrotransposons and their hosts. Cytogenet Genome Res. 2005;110: 70–90. doi:10.1159/000084940

27. Bridier-Nahmias A, Tchalikian-Cosson A, Baller JA, Menouni R, Fayol H, Flores A, et al. Retrotransposons. An RNA polymerase III subunit determines sites of retrotransposon integration. Science. 2015;348: 585–588. doi:10.1126/science.1259114

28. Patterson K, Shavarebi F, Magnan C, Chang I, Qi X, Baldi P, et al. Local features determine Ty3 targeting frequency at RNA polymerase III transcription start sites. Genome Res. 2019;29: 1298–1309. doi:10.1101/gr.240861.118

29. Carr M, Bensasson D, Bergman CM. Evolutionary genomics of transposable elements in Saccharomyces cerevisiae. PLoS ONE. 2012;7: e50978. doi:10.1371/journal.pone.0050978

30. Bleykasten-Grosshans C, Friedrich A, Schacherer J. Genome-wide analysis of intraspecific transposon diversity in yeast. BMC Genomics. 2013;14: 399. doi:10.1186/1471-2164-14-399

31. Czaja W, Bensasson D, Ahn HW, Garfinkel DJ, Bergman CM. Evolution of Ty1 copy number control in yeast by horizontal transfer and recombination. PLoS Genet. 2020;16: e1008632. doi:10.1371/journal.pgen.1008632

32. D’Angiolo M, Chiara MD, Yue J-X, Irizar A, Stenberg S, Persson K, et al. A yeast living ancestor reveals the origin of genomic introgressions. Nature. 2020;587: 420–425. doi:10.1038/s41586-020-2889-1

33. Kim JM, Vanguri S, Boeke JD, Gabriel A, Voytas DF. Transposable elements and genome organization: a comprehensive survey of retrotransposons revealed by the complete Saccharomyces cerevisiae genome sequence. Genome Res. 1998;8: 464–478. doi:10.1101/gr.8.5.464

34. Jordan IK, McDonald JF. Evidence for the Role of Recombination in the Regulatory Evolution of Saccharomyces cerevisiae Ty Elements. J Mol Evol. 1998;47: 14–20. doi:10.1007/PL00006358

35. Istace B, Friedrich A, d’Agata L, Faye S, Payen E, Beluche O, et al. de novo assembly and population genomic survey of natural yeast isolates with the Oxford Nanopore MinION sequencer. Gigascience. 2017;6: 1–13. doi:10.1093/gigascience/giw018

36. Gabriel A, Dapprich J, Kunkel M, Gresham D, Pratt SC, Dunham MJ. Global mapping of transposon location. PLoS Genet. 2006;2: e212. doi:10.1371/journal.pgen.0020212

37. Liti G, Peruffo A, James SA, Roberts IN, Louis EJ. Inferences of evolutionary relationships from a population survey of LTR-retrotransposons and telomeric-associated sequences in the Saccharomyces sensu stricto complex. Yeast. 2005;22: 177–192. doi:10.1002/yea.1200

38. Bendixsen DP, Gettle N, Gilchrist C, Zhang Z, Stelkens R. Genomic evidence of an ancient East Asian divergence event in wild Saccharomyces cerevisiae. Genome Biology and Evolution. 2021 [cited 15 Jan 2021]. doi:10.1093/gbe/evab001

39. Curcio MJ, Garfinkel DJ. Single-step selection for Ty1 element retrotransposition. Proc Natl Acad Sci U S A. 1991;88: 936–940. doi:10.1073/pnas.88.3.936

40. Lee BS, Lichtenstein CP, Faiola B, Rinckel LA, Wysock W, Curcio MJ, et al. Posttranslational inhibition of Ty1 retrotransposition by nucleotide excision repair/transcription factor TFIIH subunits Ssl2p and Rad3p. Genetics. 1998;148: 1743–1761.

41. Fournier T, Abou Saada O, Hou J, Peter J, Caudal E, Schacherer J. Extensive impact of low-frequency variants on the phenotypic landscape at population-scale. Elife. 2019;8. doi:10.7554/eLife.49258

42. Elder RT, John TPS, Stinchcomb DT, Davis RW. Studies on the Transposable Element Ty1 of Yeast I. RNA Homologous to Ty1. Cold Spring Harb Symp Quant Biol. 1981;45: 581–591. doi:10.1101/SQB.1981.045.01.075

43. Morillon A, Springer M, Lesage P. Activation of the Kss1 invasive-filamentous growth pathway induces Ty1 transcription and retrotransposition in Saccharomyces cerevisiae. Mol Cell Biol. 2000;20: 5766–5776. doi:10.1128/mcb.20.15.5766-5776.2000

44. Garfinkel DJ, Nyswaner KM, Stefanisko KM, Chang C, Moore SP. Ty1 Copy Number Dynamics in Saccharomyces. Genetics. 2005;169: 1845–1857. doi:10.1534/genetics.104.037317

45. Saha A, Mitchell JA, Nishida Y, Hildreth JE, Ariberre JA, Gilbert WV, et al. A trans-Dominant Form of Gag Restricts Ty1 Retrotransposition and Mediates Copy Number Control. Journal of Virology. 2015;89: 3922–3938. doi:10.1128/JVI.03060-14

46. Garfinkel DJ, Tucker JM, Saha A, Nishida Y, Pachulska-Wieczorek K, Błaszczyk L, et al. A self-encoded capsid derivative restricts Ty1 retrotransposition in Saccharomyces. Curr Genet. 2016;62: 321–329. doi:10.1007/s00294-015-0550-6

47. Błaszczyk L, Biesiada M, Saha A, Garfinkel DJ, Purzycka KJ. Structure of Ty1 Internally Initiated RNA Influences Restriction Factor Expression. Viruses. 2017;9. doi:10.3390/v9040074

48. Roeder GS, Fink GR. DNA rearrangements associated with a transposable element in yeast. Cell. 1980;21: 239–249. doi:10.1016/0092-8674(80)90131-2

49. Rachidi N, Barre P, Blondin B. Multiple Ty-mediated chromosomal translocations lead to karyotype changes in a wine strain of Saccharomyces cerevisiae. Mol Gen Genet. 1999;261: 841–850. doi:10.1007/s004380050028

50. Hou J, Friedrich A, de Montigny J, Schacherer J. Chromosomal rearrangements as a major mechanism in the onset of reproductive isolation in Saccharomyces cerevisiae. Curr Biol. 2014;24: 1153–1159. doi:10.1016/j.cub.2014.03.063

51. Hu WS, Temin HM. Genetic consequences of packaging two RNA genomes in one retroviral particle: pseudodiploidy and high rate of genetic recombination. Proc Natl Acad Sci U S A. 1990;87: 1556–1560. doi:10.1073/pnas.87.4.1556

52. Negroni M, Ricchetti M, Nouvel P, Buc H. Homologous recombination promoted by reverse transcriptase during copying of two distinct RNA templates. PNAS. 1995;92: 6971–6975. doi:10.1073/pnas.92.15.6971

53. Anderson JA, Teufel RJ, Yin PD, Hu W-S. Correlated Template-Switching Events during Minus-Strand DNA Synthesis: a Mechanism for High Negative Interference during Retroviral Recombination. Journal of Virology. 1998;72: 1186–1194. doi:10.1128/JVI.72.2.1186-1194.1998

54. Bleykasten-Grosshans C, Jung PP, Fritsch ES, Potier S, de Montigny J, Souciet J-L. The Ty1 LTR-retrotransposon population in Saccharomyces cerevisiae genome: dynamics and sequence variations during mobility. FEMS Yeast Res. 2011;11: 334–344. doi:10.1111/j.1567-1364.2011.00721.x

55. Mérel V, Boulesteix M, Fablet M, Vieira C. Transposable elements in Drosophila. Mobile DNA. 2020;11: 23. doi:10.1186/s13100-020-00213-z

56. McClintock B. The significance of responses of the genome to challenge. Science. 1984;226: 792–801. doi:10.1126/science.15739260

57. Hénault M, Marsit S, Charron G, Landry CR. The effect of hybridization on transposable element accumulation in an undomesticated fungal species. Verstrepen KJ, Wittkopp PJ, Bensasson D, editors. eLife. 2020;9: e60474. doi:10.7554/eLife.60474

58. Smukowski Heil C, Patterson K, Shang-Mei Hickey A, Alcantara E, Dunham MJ. Transposable element mobilization in interspecific yeast hybrids. Genome Biology and Evolution. 2021 [cited 18 Feb 2021]. doi:10.1093/gbe/evab033

59. Kawakami T, Dhakal P, Katterhenry AN, Heatherington CA, Ungerer MC. Transposable mElement Proliferation and Genome Expansion Are Rare in Contemporary Sunflower Hybrid Populations Despite Widespread Transcriptional Activity of LTR Retrotransposons. Genome Biol Evol. 2011;3: 156–167. doi:10.1093/gbe/evr005

60. Heyduk K, McAssey EV, Grimwood J, Shu S, Schmutz J, McKain MR, et al. Hybridization History and Repetitive Element Content in the Genome of a Homoploid Hybrid, Yucca gloriosa (Asparagaceae). Front Plant Sci. 2021;11. doi:10.3389/fpls.2020.573767

61. Coyne JA. Mutation rates in hybrids between sibling species of Drosophila. Heredity. 1989;63: 155–162. doi:10.1038/hdy.1989.87

62. Vela D, Fontdevila A, Vieira C, García Guerreiro MP. A genome-wide survey of genetic instability by transposition in Drosophila hybrids. PLoS One. 2014;9: e88992. doi:10.1371/journal.pone.0088992

63. Gabaldón T. Patterns and impacts of nonvertical evolution in eukaryotes: a paradigm shift. Ann N Y Acad Sci. 2020;1476: 78–92. doi: 10.1111/nyas.14471

64. Gabaldón T. Hybridization and the origin of new yeast lineages. FEMS Yeast Research. 2020;20. doi: 10.1093/femsyr/foaa040

65. Mallet J, Besansky N, Hahn MW. How reticulated are species? Bioessays. 2016;38: 140–149. doi:10.1002/bies.201500149

66. Pryszcz LP, Németh T, Saus E, Ksiezopolska E, Heged?sová E, Nosek J, et al. The Genomic Aftermath of Hybridization in the Opportunistic Pathogen Candida metapsilosis. PLoS Genet. 2015;11: e1005626. doi:10.1371/journal.pgen.1005626

67. Li H, Durbin R. Fast and accurate short read alignment with Burrows-Wheeler transform. Bioinformatics. 2009;25: 1754–1760. doi:10.1093/bioinformatics/btp324

68. Li H, Handsaker B, Wysoker A, Fennell T, Ruan J, Homer N, et al. The Sequence Alignment/Map format and SAMtools. Bioinformatics. 2009;25: 2078–2079. doi:10.1093/bioinformatics/btp352

69. Paquin CE, Williamson VM. Temperature Effects on the Rate of Ty Transposition. Science. 1984;226: 53–55. doi:10.1126/science.226.4670.53

70. Lawler JF, Haeusser DP, Dull A, Boeke JD, Keeney JB. Ty1 defect in proteolysis at high temperature. J Virol. 2002;76: 4233–4240. doi:10.1128/jvi.76.9.4233-4240.2002

